# Cerebral plasticity after hypoglosso-facial anastomosis in facial palsy: a magnetoencephalography study

**DOI:** 10.1101/2025.05.13.652175

**Authors:** Rémi Hervochon, Deborah Ziri, Guillaume Dupuch, Maximilien Chaumon, Claire Foirest, Denis Schwartz, Christophe Gitton, Nathalie George, Frédéric Tankere

**Affiliations:** Sorbonne Université, Institut du Cerveau – Paris Brain Institute – ICM, Inserm, CNRS, APHP, Hôpital de la Pitié Salpêtrière, Paris, France; AP-HP, Hôpital de la Pitié Salpêtrière, Oto-Rhino-Laryngology and Cervico-Facial Surgery Department, 75013 Paris, France; Institut du Cerveau, ICM, Inserm U 1127, CNRS UMR 7225, Sorbonne Université, CENIR, Centre MEG-EEG, F-75013, Paris, France; American Hospital of Paris, 55 boulevard du chateau, 92200 Neuilly sur Seine

**Keywords:** Facial palsy, magnetoencephalography, somatotopy, smile, hypoglosso-facial anastomosis

## Abstract

**Background.:** Hypoglosso-facial anastomosis (HFA) consists in suturing the proximal part of the hypoglossal nerve with the distal part of the facial nerve in patients with facial palsy. Axonal regrowth through the anastomosis makes it possible to restore facial motor skills, which become spontaneous after physiotherapy. This suggests cerebral plasticity.

**Objective:** We used magnetoencephalography (MEG) in a pilot study to test this hypothesis

**Methods:** Twenty-one healthy volunteers (CTRL) and 12 patients after HFA performed 5 motor tasks with MEG and electromyographic recordings: eyelid closure, smile, tongue protraction, mastication and thumb flexion. For each task, we picked the location of the maximum source activity within the precentral gyrus. We calculated the distances between this location and the vertex for each task and a somatotopy index.

**Results:** There was an interaction between the participant’s group and the task (F(4,124)=4.07, p=0.0039). In CTRL, the maximum source location was statistically different between smile and tongue tasks and between eyelid and tongue tasks (p<0.001). No such difference was observed in HFA (p=1.000). 90.5% of CTRL and 41.7% of HFA showed a normal somatotopy (p=0.0046).

**Conclusions:** In CTRL, the organization of the cortical motor areas was similar to that of Penfield’s motor Homunculus. In contrast, in HFA, eyelid closure, tongue protraction and smile areas were not significantly distinct. This supports the hypothesis of cerebral plasticity after HFA.

The Ethical Committee of Paris Idf VI approved the study (CPP Ouest 6-CPP975-HPS2).

**Financial Disclosure Statement:**

– This research has received funding from the “**Fondation pour la Recherche Médicale**” and from the “**Fondation des Gueules Cassées**”. It was performed on a platform of the France Life Imaging network partly funded by the grant “ANR-11-INBS-0006” and by the program “Investissements d’avenir” ANR-10-IAIHU-06. The funding bodies had no role in the study design, the data collection, analysis or interpretation, or the article writing.
– The authors have neither commercial association nor financial disclosure.

Generative AI was not used

## 1. INTRODUCTION

In case of severe and permanent peripheral facial palsy (PFP), patients may benefit from surgical rehabilitation procedures in order to reanimate partially the paralyzed side. Hypoglosso-facial anastomosis (HFA) consists in suturing the proximal part of the hypoglossal nerve with the distal part of the facial nerve (Lamas et al., 2015; Sargent, 1912). Axonal regrowth through the anastomosis makes it possible to obtain facial motor skills after 6 months of specific physiotherapy. It allows obtaining a voluntary and even a spontaneous smile. This suggests cerebral plasticity. However, this phenomenon is yet to be demonstrated and fully understood. A better understanding of plastic processes should allow us to validate or modify our postoperative physiotherapy and speech therapy techniques in order to optimize the functional results.

Magnetoencephalography (MEG) is a functional brain imaging method that allows the non-invasive detection of the magnetic fields produced by the electrical activity of neuronal ensembles. It has been used to characterize the somatotopic organisation of the facial sensorimotor cortex(Nakahara et al., 2004; Nakamura et al., 1998; Spooner et al., 2022; Yamashita et al., 1999; Yang et al., 1993). The identification of purely sensory cortical areas of the face and the tongue has already been conducted with MEG in healthy subjects for different types of stimuli (tactile, electrical, vibratory) (Karhu et al., 1991; Yamashita et al., 1999). Face motor functions were also studied, during speech tasks (Memarian et al., 2012), and motor tasks, such as tongue protrusion (Maezawa et al., 2017, 2014; Nakasato et al., 2001). MEG—coupled with magnetic resonance imaging (MRI) for the localization of the brain sources of magnetic activities—allowed the precise identification of the cortical motor areas of face, tongue, and hand in healthy subjects (Cheyne et al., 2006; Kirsch et al., 2007). For example, for finger movements, a motor magnetic field was localized in the contralateral precentral gyrus. Its maximum amplitude was observed about 50ms before the onset of the movement (Cheyne et al., 2006). The magnetic evoked fields time-locked to movement and measured prior to it reflect brain activities related to motor control; they are particularly interesting to study because they are not contaminated by muscular contraction artefacts. The cortical activation prior to movement has been studied using source localization of neuromagnetic activities and it allowed to detect low signal-to-noise ratio activities (Cheyne et al., 2006; Maezawa et al., 2014).

Here, we performed a pilot study with MEG to map the somatotopy of the face in controls and in patients with facial palsy rehabilitated by HFA, during simple motor tasks of the face and of the thumb. Our main objective was to obtain preliminary evidence for the face motor cortex reorganization in patients operated by HFA. Normal somatotopy of motor representations in the precentral gyrus regions of the motor cortex is well established (Becker, 1953; Grabski et al., 2012). The motor representations of body parts are organized along the precentral gyrus convexity as follows: leg, arm, hand, face, and swallowing are represented in this order from the vertex to the lateral fissure. Face representation is localized in the middle frontal part of the precentral gyrus and it is also somatotopic with eyelids, mouth, tongue, and mastication represented in this order from top to bottom along the gyrus (Figure 1).

**Figure 1:**
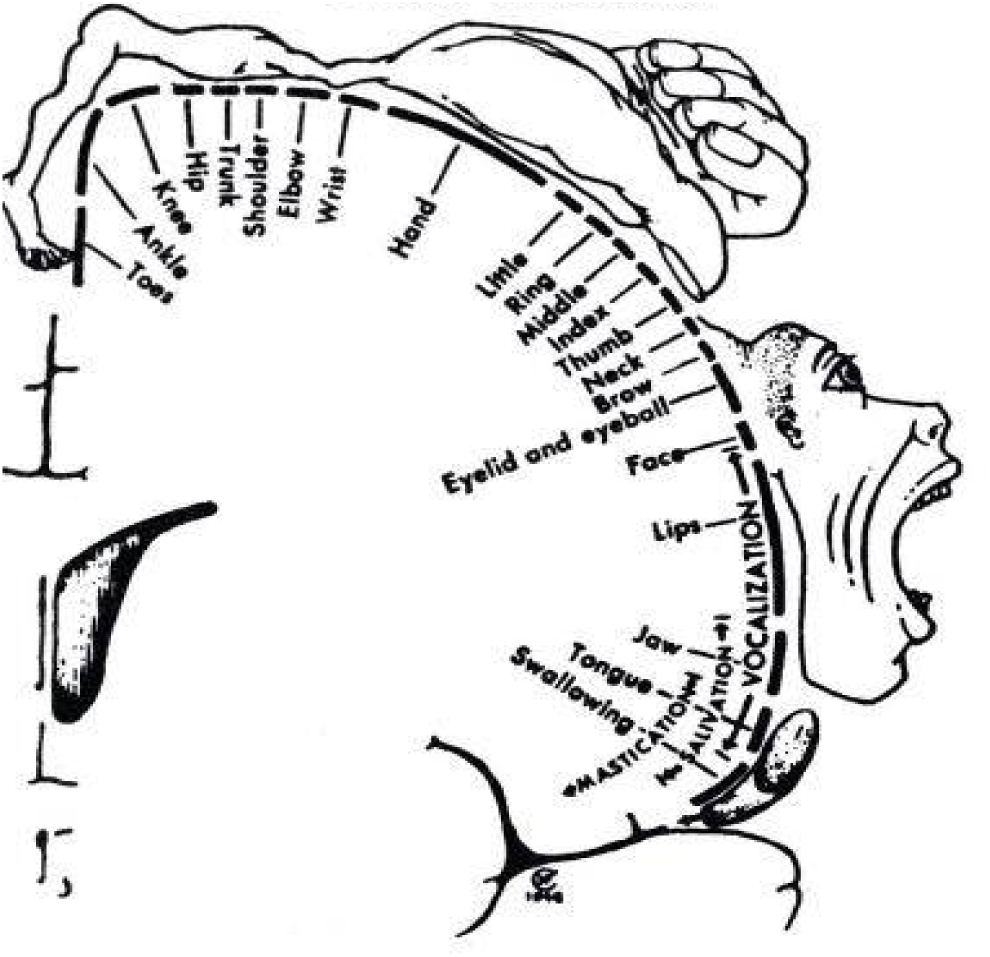
Normal motor Homunculus.

We expected to be able to map face somatotopy by localizing the sources of magnetic fields evoked prior to simple movements of different face parts in controls and in patients operated by HFA. Our hypothesis was that whereas controls would have a normal somatotopy, there would be changes in the somatotopic organisation of face representation after HFA, particularly concerning the representation of smile, eyelid closure, and lingual protraction.

## 2. MATERIALS AND METHODS

### 2.1. Participants

Twenty-one healthy volunteers (CTRL) (13 men, mean age ± SD = 40.6±13.5 years) and 12 patients (8 men, mean age ± SD = 45.8±12.1 years) with facial palsy who had underwent HFA were included. The CTRL and HFA groups did not differ either in male/female sex ratio (Fisher’s exact test, p=1) or in age (unequal variance Student’s test, p=0.25; Table 1). All patients underwent surgery in the Oto-Rhino-Laryngology and Cervico-Facial Surgery department in Pitié-Salpêtrière Hospital in Paris, France. Patients who had undergone successful HFA surgery for more than 6 months, who were not lost to follow-up, and who met the inclusion and exclusion criteria were offered to participate in the study. The delay between the surgery and MEG recording session was between 9 and 90 months (Table 1). All subjects (CTRL and HFA) were right-handed. The Ethical Committee of Paris Idf VI approved the study (CPP Ouest 6-CPP975-HPS2). Exclusion criteria were: being under 18 years old, having an MRI contra-indication, presenting a progressive neurological or psychiatric disease, taking drug with known action on the central nervous system.

**Table 1:** Demographic data of patients and controls.

|  | Sex (man=1) | Age (years) | Delay HFA-MEG (months) |
| --- | --- | --- | --- |
| HFA1 | 0 | 37 | 16 |
| HFA2 | 1 | 46 | 9 |
| HFA3 | 1 | 36 | 70 |
| HFA4 | 1 | 54 | 54 |
| HFA5 | 0 | 55 | 48 |
| HFA6 | 1 | 37 | 48 |
| HFA7 | 0 | 30 | 19 |
| HFA8 | 1 | 29 | 77 |
| HFA9 | 1 | 56 | 27 |
| HFA10 | 1 | 51 | 62 |
| HFA11 | 0 | 49 | 90 |
| HFA12 | 1 | 69 | 32 |
| CTRL1 | 1 | 65 |  |
| CTRL2 | 1 | 26 |  |
| CTRL3 | 0 | 28 |  |
| CTRL4 | 1 | 29 |  |
| CTRL5 | 0 | 54 |  |
| CTRL6 | 0 | 51 |  |
| CTRL7 | 0 | 47 |  |
| CTRL8 | 0 | 48 |  |
| CTRL9 | 1 | 22 |  |
| CTRL10 | 1 | 38 |  |
| CTRL11 | 1 | 48 |  |
| CTRL12 | 0 | 24 |  |
| CTRL13 | 0 | 46 |  |
| CTRL14 | 1 | 27 |  |
| CTRL15 | 1 | 52 |  |
| CTRL16 | 1 | 48 |  |
| CTRL17 | 1 | 42 |  |
| CTRL18 | 0 | 23 |  |
| CTRL19 | 1 | 39 |  |
| CTRL20 | 1 | 29 |  |
| CTRL21 | 1 | 66 |  |

### 2.2. Apparatus and data acquisition

All participants underwent MEG recordings and anatomical T1 MRI scanning at the Centre de Neuroimagerie de Recherche (CENIR) of the Paris Brain Institute in Paris.

#### 2.2.1 MEG

During MEG recordings, participants were seated in a dimly lit, magnetically shielded room, with a screen display placed at a viewing distance of 85 cm. Instructions were delivered visually on this screen.

Neuromagnetic fields were recorded with a 306-channels whole-head Elekta Neuromag® Triux MEG system comprising 204 orthogonally oriented planar gradiometers and 102 radial magnetometers regularly distributed at 102 locations over the scalp. Magnetic signals were recorded continuously with a sampling rate of 1 kHz and a low-pass filter set at 330Hz.

Four head position indicator (HPI) coils were affixed to the subject’s head. The positions of the HPI coils and of multiple points on the scalp were digitized with a magnetic digitizer (Polhemus 3D Fastrack system). These positions were recorded with respect to a head coordinate frame defined by three anatomical landmarks: the nasion and the left and right tragi. These points were later used to coregister the coordinate systems of the head, the MEG sensors, and the subject’s anatomical MRI. For eye movements and blinks, standard electrooculograms (EOG) were recorded by disposable bipolar electrodes, with two electrodes placed above and below the left eye respectively. The electrocardiogram (EKG) was monitored by two disposable electrodes placed on the left belly and the right collarbone.

Electromyograms (EMG) were recorded with pairs of reusable electrodes placed on the skin above the following facial muscles: temporalis (used for mastication), orbicularis oculi (used for palpebral occlusion, that is, eye closing), zygomaticus (used for smiling), genioglossus (used for tongue protraction); in addition, two electrodes were placed on the flexor pollicis brevis (used for thumb flexion). The EMG, EKG, EOG, and MEG signals were recorded simultaneously and synchronously at the sampling frequency of the MEG system (1kHz). (Figure 2).

**Figure 2:** Placement of electromyographic electrodes used for the recording of the temporalis, orbicularis oculi, zygomaticus, and genioglossus.

#### 2.2.2. Anatomical MRI

A T1-weighted morphological MRI was acquired in all subjects after the MEG recordings. The MRI scanner was a Siemens 3 Tesla PRISMA with a 64-channels antenna. The sequence parameters were: TR=2400.0ms, TE=2.22ms, TI=1000ms, sagittal acquisition, 0.8mm×0.8mm×0.8mm voxel size, 256mm×240mm×204.8mm acquisition field (i.e. 320×300×256 acquisition matrix).

### 2.3. Task and procedure

The participants performed 5 blocks of motor tasks. Each block was composed of 5 motor tasks: “thumb flexion” (Th), “eyelid closure” (E), “smile” (S), “tongue protraction” (To), and “mastication” (M). Each task was performed repeatedly for 48 seconds. The tasks were performed in a different, randomized order in each block. The task instructions were presented on screen for 5 seconds; it was then replaced by a fixation cross, which indicated to the subject to start performing the requested movement. The subject repeated the movement, self-paced at around 1Hz, for 48 seconds, until a “stop” instruction was displayed. Then, a new instruction was displayed after about 10 seconds had elapsed. Thumb flexion was performed with the right hand by the healthy volunteers and the patients operated on the right side and with the left hand by the patients operated on the left side. At the end of the motor task blocks, we recorded a 5-minutes resting state. The subject had to remain still with eyes opened, fixating a point at the centre of the screen.

### 2.4. Data processing

#### 2.4.1. EMG data

EMG data were used to mark the onset of each movement. These onsets were marked manually on the recordings as the moment when the EMG signal deviated by more than ∼1µV from the level it had during the preceding 100ms. The signal was then epoched from –500ms to +200ms around the EMG marker. Trials with blinks within 500ms before movement onset were excluded. The average number of valid motor events per subject, without any artefact, was 124.9±88.1 for (Th), 106.7±49.3 for (E), 82.3±46.1 for (S), 85.2±40.2 for (To), and 85.0±40.2 for (M).

#### 2.4.2. MEG data

Denoising of the raw MEG data was done with *Maxfilter* (version 2.2.10; Elekta AB, Stockholm, Sweden) using temporal signal space separation (tSSS) to remove interferences from magnetic fields originating outside of a sphere delimited by the sensor helmet. The magnetic fields time-locked to movement onset were then averaged across trials for each type of movement and each subject.

The resting state data were screened for muscular artefacts or data segments with signal amplitude larger than 4 standard deviations (SD) above the mean signal. Independent Component Analysis (ICA) was used to remove cardiac and ocular artefacts; it was performed with the extended infomax algorithm implemented in Fieldtrip (Oostenveld et al., 2011).

Subsequent data processing and analysis were performed using *Brainstorm* (version of 07/15/2017), which is documented and freely available for download online under the GNU general public license (http://neuroimage.usc.edu/brainstorm) (Tadel et al., 2011).

#### 2.4.3. Source localization

Source localization was performed using a weighted minimum norm estimate (wMNE) algorithm. First, for each subject, the anatomical MRI was segmented with the Freesurfer image analysis suite. We used the surface defined by the cerebrospinal fluid – grey matter interface as our source space: the original mesh was subsampled with 15,000 vertices. A source was placed at each vertex with an orientation normal to the local surface. Then, we computed the forward model using overlapping spheres. The forward model (or so-called head model in Brainstorm) allows modelling how neural electric currents from the sources produce the magnetic fields at the sensor level, taking into account the geometry and electrical properties of head tissues.

The resting state data were used to compute the noise covariance matrix for each subject. Source amplitudes across time were computed for each trial thanks to the inverse operator computed from the noise covariance matrix and the forward operator. Source amplitudes were then averaged across trials for each type of movement. Baseline activity was subtracted using the mean source amplitude between –500 and –400ms before movement onset. Source amplitudes were then taken as the absolute value of the baseline-corrected activities. Finally, for group-level analysis, sources estimated in each task for each subject were projected on a reference brain model (MNI152) (Evans et al., 1994).

#### 2.4.4. Source analysis

We computed the source average amplitudes in the [-100ms; –50ms] time window before movement onset for each subject and each task. We localized manually the maximal source, that is, the point with maximal amplitude, within the left precentral gyrus (delineated according to the Desikan-Killiany atlas (Desikan et al., 2006)) in the healthy volunteers and in the patients operated on the right side, and within the right precentral gyrus in the patients operated on the left side. We made this choice because the hypoglossal nucleus receives afferents from the contralateral corticonuclear tract(Simon and Mertens, 2009). We noted the coordinates of the maximal source in the MNI space (X,Y,Z). We measured the distance between the maximal source and a reference point (X_0_,Y_0_,Z_0_) defined as the top of the precentral gyrus (point with largest Z coordinate along the precentral gyrus). For this, we first computed the Euclidean distance l between the identified maximal sources and the top of precentral gyrus, as follows:

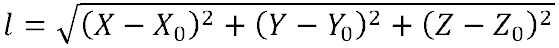

Then, we transformed this distance *l* into the length *L* of an arc, considering the precentral gyrus coronal convexity as the arc of a circle with radius *r* = 64.35 mm; this radius corresponded to half the cross-sectional diameter of the average brain. We used the following formula:

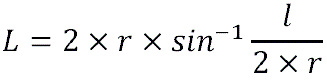

This was taken as the estimated position of the source within the precentral gyrus for each type of movement and for each subject.

### 2.5. Statistical analysis

The statistical analyses were performed using the freely available JASP software (Version 0.18.3; University of Amsterdam; https://jasp-stats.org/).

We tested for differences in the source localization of the 5 types of movements. For this, we performed an ANOVA with the L distance as the dependent variable, group (CTRL / HFA) as between-subject factor, and task (Th, E, S, To and M) as within-subject factor. We used Greenhouse-Geisser correction in order to correct for departures from sphericity assumption in the task factor that had 5 levels. We also performed 2-by-2 between-task post-hoc tests, focusing on the following comparisons of L distances within each group: E vs. To, S vs. To. For this, we used t-tests with Holm correction for multiple comparisons, as implemented in JASP. In addition, we used a two one-sided test (TOST)(Schuirmann, 1987) procedure to test for equivalence in L distance between E and To tasks and between S and To tasks in the HFA group, using the TOSTER package of R software(Lakens, 2017).

We used a somatotopy index (SI) inspired by Meunier et al. (Meunier et al., 2001). This measure assumes that the source peaks along the precentral gyrus should follow the order corresponding to the known somatotopy of motor representations. It measures deviations from this order with the following formula:

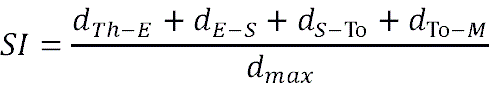

Where *d_X-Y_* is the distance between source peaks for tasks X and Y (tasks being: Th, E, S, To, and M), and d_max_ is the distance between the two most distant source peaks for a given subject. Tasks are expected to be found in the order Th, E, S, To, M, along the precentral gyrus, following Penfield’s homunculus. The SI is therefore either equal to 1, in the case where cerebral sources are found in the exact order defined above, or different from 1, in the case where the peak sources are in a abnormal order. A Fisher’s exact test was used to compare the proportion of subjects with a SI equal to 1 between the CTRL and HFA groups.

## 3. RESULTS

The L distances for the maximum source of each task in both the HFA and the CTRL groups are presented in Figure 3.

**Figure 3:**
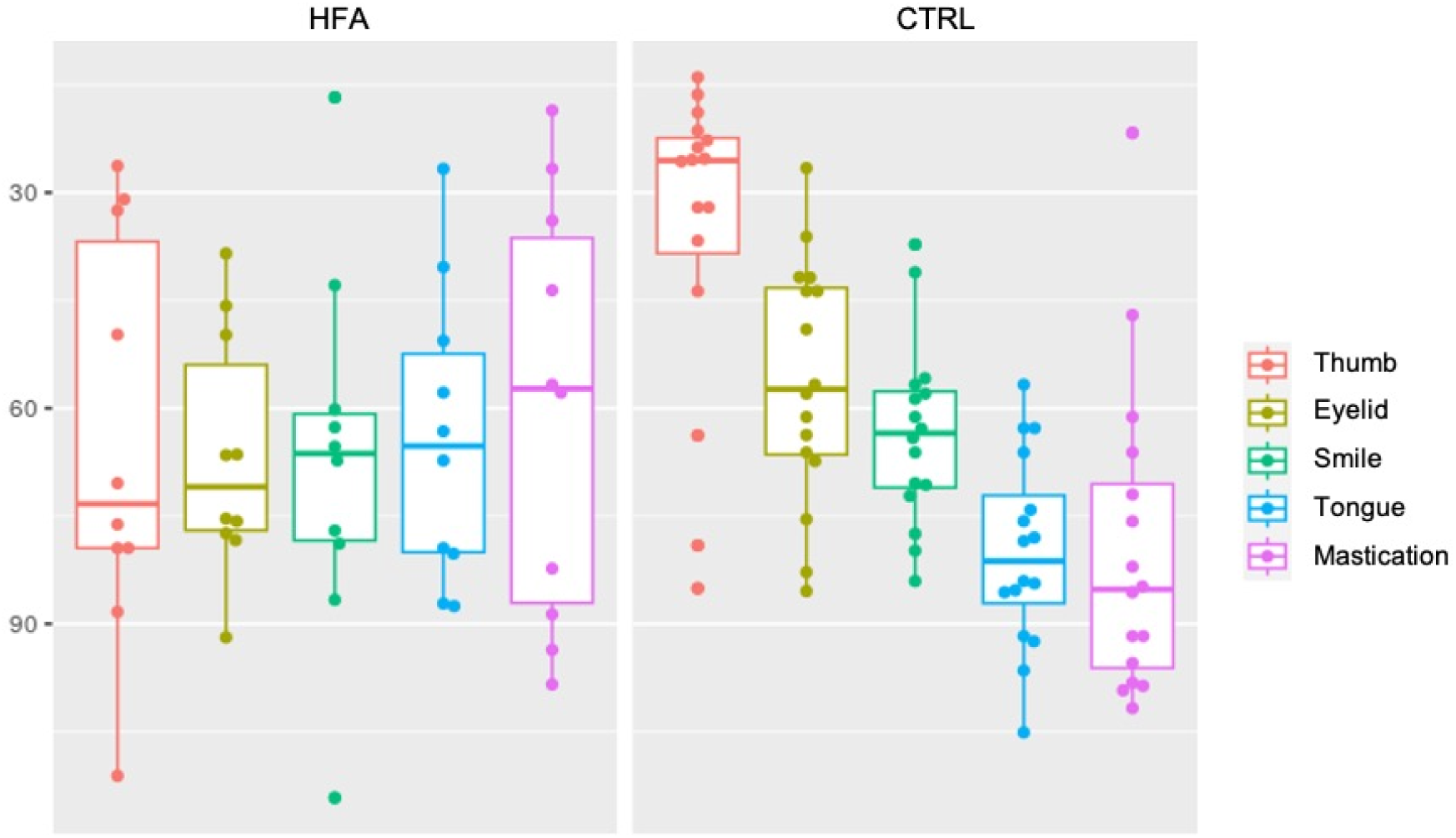
Distance L for each subject and each motor task. Each point is one subject in a given task. Standard box plots are used. The horizontal line in the box represents the median.

In CTRL, the L distances (mean±SD) were 29.2±11.9 mm, 54.0±12.7 mm, 65.4±12.8 mm, 83.7±13.4 mm and 88.1±19.9 mm for the Th, E, S, To, and M tasks, respectively, which corresponded to the normal somatotopy. In HFA, the L distances were respectively 47.0±22.8 mm, 65.5±15.5 mm, 66.7±20.2 mm, 72.3±23.3 mm and 76.8±32.1mm (Figure 3).

The ANOVA revealed a marked effect of the task (F(4,124)=28.59, LJ_GG_=0.783, p<0.001). There was no significant main effect of the group (F(1,31)=0,30, p=0.59). The interaction between group and task was statistically significant (F(4,124)=4.07, LJ_GG_=0.783, p=0.008).

For the CTRL group, there were statistically significant differences between the eyelid closure and the tongue protraction tasks and between the tongue protraction and the smile tasks (Table 2). In contrast, no such difference was observed in HFA (both t<1). We further performed equivalence tests (TOST) between these tasks in the HFA group, with boundary for equivalence set at 10% (7.5 mm). It did not yield any significant result (Table 2).

**Table 2:** Post-hoc tests: comparison of interest. IC95%, corrected t-test and p-value.

|  | CTRL | HFA |  |
| --- | --- | --- | --- |
|  | Student | Student | TOST |
| Eyelid - Tongue | [-42.669 ; -16.589] | [-34.573 ; 21.101] | t(11) = 0.962 |
|  | t = 6.560 | t = 0.715 | p = 0.822 |
|  | p < 0.001* | p = 1.000 |  |
| Smile - Tongue | [-31.288 ; -5.210] | [-33.370 ; 22.304] | t(11) = 1.622 |
|  | t = 4.040 | t = 0.587 | p = 0.933 |
|  | p < 0.001* | p = 1.000 |  |

In the CTRL group, 19 out of the 21 participants (90.5%) had a SI=1. By contrast, in the HFA group, only 5 out of the 12 patients (41.7%) had a SI=1. Fischer’s exact test revealed a significant difference between the two groups (p=0.0046).

## 4. DISCUSSION

The cortical motor areas dedicated to the thumb, eye closure, smile, lingual protraction and mastication were organized, as expected, according to Penfield’s motor homunculus (Becker, 1953) in the healthy volunteers, underlying the value of MEG for cortical source localization. The coordinates of maximal source activity found here are similar to those described by Grabski for the lips and tongue movements (Grabski et al., 2012), and those identified by our team (Hervochon et al., 2024). By contrast, in the patients operated with HFA, the organization was different, with altered face somatotopy in comparison to controls. In particulier, the peak sources of the three motor functions concerned by HFA (eyelid, smile and tongue tasks) seemed to overlap (Figure 3) and their localisation was not statistically different.

Normal motor brain mapping has been shown in functional imaging in many cases. Although the description of the homunculus is relatively old (Becker, 1953; Brodmann, 1909; Geyer et al., 2011), some recent works confirmed it with fMRI (Martin et al., 2004) (Romeo et al., 2013). As suggested by Takai et al., it corresponds to “somatotopy with overlap” (Takai et al., 2010). On the other hand, Gordon has recently completely redesigned and challenged the dogma of the homunculus: Using fMRI, they discovered that the classic homunculus is interrupted by regions with sharply distinct connectivity, structure, and function, alternating with effector-specific (foot, hand, mouth) areas. The inter-effector regions lacked movement specificity and co-activated during action planning. Thus, two parallel systems are intertwined in motor cortex to form an integrate-isolate pattern: effector-specific regions (foot, hand, mouth), and a mind-body interface for the integrative whole-organism coordination (Gordon et al., 2022).

Cerebral plasticity is the capacity of the brain to modify its functional organization after pathological transformations of the central nervous system, during learning, during environmental interactions, or after modification of body structures (blindness, amputation, surgery). Plasticity is maximal in childhood but also occurs in adults (Hansson and Brismar, 2003). This plasticity can take the form of a variation in cerebral activity intensity, of an extension of the activity next to the functional area, or of the solicitation of new regions far from the original functional area.

Brain plasticity has been studied after other surgical rehabilitations of facial palsy. Garmi et al. used fMRI and showed an enlargement of the smile area, which merged with the mastication area, in 10 patients at 3 months and 1 year after Lengthening Temporalis Myoplasty (LTM) surgery (Garmi et al., 2013). Hervochon et al. showed with MEG a convergence of the peak sources for smile and mastication in a LTM group (Hervochon et al., 2024). Interestingly, another study by Buendia et al. with fMRI showed similar brain activity for smile and mastication in facial palsy patients rehabilitated by a transfer of the masseteric nerve, whose initial function is masticatory (Buendia et al., 2016). Taken together, these results suggest that peripheral muscle or nerve suture may lead to similar plasticity at the brain level.

To the best of our knowledge, only few studies investigated plasticity after HFA. Bitter et al. found that lip movements were associated with activation of both face and tongue cortices in seven HFA patients (Bitter et al., 2011). Willer et al., Tankéré et al. and Ling et al. showed that HFA induced central plastic changes in the blink reflex circuitry in patients (Tankéré et al., 2000; Willer et al., 2002), (Ling et al., 2021). Neuronal plasticity was also demonstrated in the rat (Neiss et al., 1992) and cat (Vera et al., 1975). Altogether, these studies and ours support the view that rehabilitative surgery of facial palsy is associated with functionally specific plasticity of the motor cortex. We did not find any statistically significant difference in terms of peak sources between eyelid closure, smile and tongue protraction task in HFA. Although the equivalence test was not significant, it seemed that the peak sources showed more overlap for these tasks in the HFA than in the CTRL groups. Indeed, Figure 3 illustrates that the distribution of motor tasks is well organized in ctrl – in agreement with Penfield’s homunculus – whereas the 3 facial motor tasks appear to overlap in HFA. This suggests functional remodelling of the motor cortex of the face following HFA: The motor area of the tongue would be solicited after HFA surgery and physiotherapy to produce facial motor skills (closing of the eyelids and smiling). The cortex functional reorganization found may suggest potential treatment targets in the central nervous system for adjuvant therapies such as repetitive transcranial magnetic stimulation to further improve functional recovery (Ling et al., 2021).

One may wonder why 41.7% HFA showed a normal somatotopy. We can hypothesize that brain plasticity occurred in our patients given that they all had functional HFA; yet, our results show that this is not necessarily observed with neuromagnetic source imaging. This suggests that MEG cannot capture all brain reorganizations. This is not surprising since we only focused on 5 simple motor tasks among an infinitely more complex and vast motor cortex. Another possibility is that there is inter-individual variation in the extent of brain plasticity following HFA, especially with heterogeneous time intervals between surgery and MEG. Future studies with a greater sample of participants could allow investigating this.

In counterpoint with our conclusions, one could hypothesize that our result does not reflect the functional remodelling of the brain, but rather the fact that in order to produce facial motor tasks, the patient must activate brain area dedicated to tongue muscles. According to the literature and our clinical experience, 6 months after surgery most patients are able to dissociate facial motor skills and tongue movement. This was the case for our 12 patients who were able to smile spontaneously. Whether they “pull out their tongue to smile” or “smile spontaneously”, our results suggest cortical changes to produce a smile or eyelid closure.

We can also question the non-pathological nature of overlapping. Indeed, the famous figure 1 and the work of Takai et al. suggest physiological overlapping in normal subjects (Takai et al., 2010). This is why we compared HFA patients to controls. Our results, although preliminary due to our small sample of participants, indicated that overlapping was greater after HFA than in controls.

MEG offers excellent temporal resolution which allowed us to focus on the milliseconds before the start of the movement and to study pre-motor brain activity. The disadvantage of MEG is poorer spatial resolution than functional MRI. However, functional MRI was impossible given the artifacts linked to facial movements and its poor temporal resolution.

Our findings should be considered as preliminary, since the small sample size of our study limits their generalizability. Our statistical power was limited and we were therefore bound to be able to detect only large effects. Consequently, the size of the effects reported here should be considered with caution; it was likely overestimated. The repeated inter-measure or intergroup differences on MEG signals are usually about 10% (or 1 standard deviation). With these conditions, the sample size used in most brain imaging studies and recommended to detect such differences is 15 to 25 (Friston, 2012). Yet, it is noteworthy that we were able to show evidence for functionally relevant brain plasticity based on MEG in a group of 12 patients after HFA compared with 21 healthy controls.

Another limitation was the absence of preoperative or longitudinal data. This raises the possibility that the changes that we observed were not solely due to HFA surgery and ensuing facial motor skills rehabilitation. Further research with larger sample size, different types of surgical procedure of smile rehabilitation, and longitudinal data will be needed to fully characterize the brain plasticity ensuing from facial palsy surgery.

Brain plasticity after peripheral surgery has been shown in other sensorimotor domains. For example, syndactyly is a pathological condition that can be treated by surgical intervention. It is a congenital malformation characterized by the fusion of two or more fingers, which can be separated surgically. Mogilner et al. showed that before surgery the cerebral representation of the hand was reduced and without any somatotopy, whereas after surgery each finger had a distinct cortical region (Mogilner et al., 1993). In this case, plasticity occurred very quickly, only a few hours after surgery (Stavrinou et al., 2007). All these studies demonstrate the great plasticity of the human brain which is able to reorganize itself after peripheral surgeries.

An important feature of MEG is that it allowed us to localize the sources of the motor command activated just before the movements. We chose to study the time window [–100msec; –50msec] for several reasons. Performing facial motor tasks generates big muscular artefacts, which make the MEG signal analysis impossible during the movement. Therefore, it is advantageous to use a time window quite far from (at the time scale of electric signal propagation) and nevertheless just before the beginning of the movement, when pre-movement motor cortex activity would be maximal. Furthermore, Cheyne et al. showed maximal pre-motor activity amplitude 50ms before the beginning of movement (Cheyne et al., 2006). This led us to choose a window of 50msec, centred on –75msec before the beginning of the movement.

An important question that remains is: Could the differences that we observed be related to facial palsy rather than to surgery? Several authors have shown, after peripheral deafferentation, a functional cerebral reorganization persisting despite good clinical recovery, and a decrease in functional connectivity (Klingner et al., 2014, 2011; Rijntjes et al., 1997; Rödel et al., 2004). Using transcranial magnetic stimulation and positron emission tomography, an increased activation of the sensory-motor cortex of the hand on the contralateral side to facial palsy was reported, with its extension to the presumed area of the face. However, Garmi et al. did not find any difference in fMRI between healthy volunteers and patients before surgery. To answer this important question, we started a prospective study to compare patients with facial palsy before versus after surgery.

## 5. CONCLUSION

In CTRL, the organization of the cortical motor areas was similar to that of the motor Homunculus of Penfield. In HFA, the peak sources were found to be not statistically different for the eyelid, smile and tongue tasks and 58.3% of patients did not display the normal somatotopy. These results were obtained in a small sample of patients and will need to be extended in the future, to assess post-versus pre-surgery somatotopy of the motor cortex on a within-subject basis. They support the view that MEG is a valuable non-invasive imaging technique for assessing brain functional organization and plasticity.

## Acknowledgments / Fundings

This research has received funding from the “**Fondation pour la Recherche Médicale**” and from “**Fondation des Gueules Cassées**” and was performed on a platform of France Life Imaging network partly funded by the grant “ANR-11-INBS-0006” and by the program “Investissements d’avenir” ANR-10-IAIHU-06. The funding bodies had no role in the study design, the data collection, analysis or interpretation, or the article writing.

## Authorship confirmation/contribution statement

Conceptualization: FT, NG, RH, DS

Data curation: RH, MC, DZ, CF, DS, CG, GD

Formal Analysis: MC, NG, RH, DZ, CF, GD

Methodology: FT, NG, RH, DS

Software: MC, DS, CG

Supervision: NG, FT, DS

Writing – original draft: RH, MC, DZ, CF, DS, CG, NG, FT, GD

Writing – review & editing: RH, MC, DZ, CF, DS, CG, NG, FT, GD

## Statements and declarations

**none**

## Ethical considerations

The Ethical Committee of Paris Idf VI approved the study (CPP Ouest 6-CPP975-HPS2).

## Consent to participate

written, approved by the IRB

## Consent for publication

informed consent for publication was provided by the participant

The author(s) declared no potential conflicts of interest with respect to the research, authorship, and/or publication of this article

